# Flight crashes have demographic consequences in a long-lived seabird

**DOI:** 10.64898/2026.01.09.698620

**Authors:** Stefan Barnett, James Bull, Andrew N Ross, Sarah Wanless, Emily LC Shepard

## Abstract

Risk is a key currency in animal ecology, yet studies of risk have almost exclusively focused on predation. Accidents, defined as a momentary loss of control, represent another potential source of injury and mortality. Such events are seldom documented and are generally assumed to be vanishingly rare in natural systems. Nonetheless, regular crash-based mortality has been documented in a population of northern gannets through monthly surveys conducted over three years in the 1970s. We revisit these data, using hindcasting, environmental records and computational fluid dynamics (CFD) models to investigate the environmental drivers of gannet crashes, and matrix population models to examine their demographic consequences under a range of environmental scenarios. Wind direction emerged as the sole predictor of crash-based mortality, with the probability of crashes increasing in north-westerly winds. CFD models revealed that north-westerlies are associated with a marked increase in turbulent kinetic energy along the breeding cliffs, relative to the other modal wind direction, which likely challenges flight control. A total of 367 crashes accounted for 5.4% of annual adult mortality. Removing this mortality led to a projected increase in population of 23.9% over 50 years, equivalent to an additional 10,583 individuals. Overall, this suggests that turbulence close to the substrate can result in fatal losses of flight control that can impact population-level processes. Our results also highlight the need for greater understanding of how airflows influence the risk of accidents across flying animals, both in current and changing wind regimes.

## Introduction

The risk of predation has a profound impact on animal ecology, impacting animal physiology (Sheriff and Thaler 2014), decisions (Hebblewhite and Merrill 2007) and space-use (Dellinger et al. 2019). Accidents, caused by momentary loss of control, can also result in injury or mortality (Wheatley et al. 2021). Animals appear to be sensitive to these risks; gulls select flight paths with weaker updrafts to reduce collision risk (Shepard, Williamson, Windsor 2016) and small arboreal mammals reduce their speed when foraging along thinner branches (Wheatley et al. 2018). Nonetheless, accidents themselves are assumed to be rare in natural systems as they are seldom documented. The potential for these risks to influence population-level processes therefore remains unknown.

Accidents are particularly pertinent for flying animals, as flight is fast - typically between 5 and 20 m/s (Hedenström 2002; Tobalske et al. 2003). Split-second losses of flight control therefore have the potential to be catastrophic when animals are operating close to solid surfaces. The risks of flight close to ground explains why > 60% of fatal accidents in aircraft occur during take-off or landing, even though these phases represent tiny proportions of the overall flight time (Boeing Company 2017). Two studies suggest that accidents can cause substantial and repeated mortality in free-flying animals. Schoombie et al. (2023) found ∼100 carcasses a year for almost 4 years in a colony of grey-headed albatross (*Thalassarche chrysostoma*), with many carcasses showing signs of injury consistent with collisions. While this study did not test whether crash frequency was predicted by wind conditions, the concentration of carcasses in an area with a down-draft close to the breeding ridge suggests that airflows compromised flight control.

Regular crash fatalities also occur in northern gannets (*Morus bassanus*) (hereafter, “gannets”) breeding on Ailsa Craig in Scotland, one of the major breeding colonies in the Northeast Atlantic (Wanless 1979). This colony is distributed along west-facing cliffs that end in talus slopes, rather than the sea. In the 1970s, Wanless (1979) found gannet carcasses along these talus slopes. Many had sustained injuries such as broken wings, beaks and necks, consistent with uncontrolled collisions with the ground. Regular counts of these carcasses revealed a strikingly high instance of crashes, with 367 carcasses recorded over three years, corresponding to one fatal crash every other day during the breeding season.

Surprisingly, crashes could not be explained by lack of flight experience as 343 of 367 carcasses were adult birds (Wanless 1979). Age therefore does not protect individuals against these accidents and the causes remain unknown. We hypothesise that the probability of crashes is related to environmental conditions, with wind and fog being the most likely predictors. Different elements of the wind could be detrimental for flight control; there are records of albatrosses sustaining fatal injuries when landing in low winds (Cone 1964), presumably as they rely on headwinds to reduce their groundspeed and landing impact, whereas downdrafts appeared to be detrimental for grey-headed albatrosses (Schoombie et al. 2023). Alternatively, birds may be impacted by poor visibility i.e. sea fog, which has been linked to grounding in shearwaters (Syposz et al. 2018). Finally, biological factors could increase the likelihood of crashes, if landing becomes challenging due to crowding at the colony, which changes through the breeding season (Schippers et al. 2011).

Crash-based mortality only represents 0.5 % of the overall gannet population breeding on Ailsa Craig (Wanless 1979). Nonetheless, gannet populations, like other long-lived species, are sensitive to even small changes in adult mortality due to their naturally high adult survival (0.922, Wanless et al. 2006) and longevity (adults can reach at least 37 years (British Trust for Ornithology 2010). The high number of adult fatalities on Ailsa Craig therefore means that accidents are likely to have demographic consequences. In this study, we investigate whether environmental factors influence individual crash risk in gannets breeding on Ailsa Craig and whether this, in turn, could impact population growth rates under both current and future environmental conditions. We address the following specific objectives: (1) investigate whether fatalities could be predicted by environmental factors, using statistical and computational fluid dynamics (CFD) models to assess the role of wind and fog. We then (2) quantify the impact of crash mortality on the population growth rate, using matrix population models (MPMs) to project population trajectories from stage-specific survival transitions (Logofet 2002), and predicting that even a small number of adult crashes could have a significant effect on the population trajectory (Maestri et al. 2017; Spencer and Janzen 2010). Finally, we (3) integrate outputs from the statistical models with the MPMs to predict how changing wind conditions would influence gannet population trajectories. Together, this should provide mechanistic insight into the drivers of accidents in flying animals and the potential for accidents to affect population-level processes.

## Methods

### Carcass data

Carcass data from Ailsa Craig were based on monthly surveys conducted between February and October from 1974 to 1976 (Wanless 1979). Surveys were conducted along the talus slopes that lie underneath many of the of the breeding cliffs on the south-western side of the island (Wanless 1979) (table 1). Carcasses were aged as adult or immature based on plumage. Most showed signs of traumatic injury, such as broken wings, necks, or beaks, as well as internal haemorrhaging suggesting collision as the cause of death. The counts represent a minimum level of crash-based mortality, as the talus slopes are steep and covered in dense vegetation, creating challenging survey conditions. Carcass numbers were used in objective 1, where they were modelled in relation to environmental conditions and objective 2, where they were used to assess the demographic consequences of removing crash-based mortality.

**Table 1.** Carcass count data used in MPMs to adjust transition probabilities as well as statistical models. Collected at the base of Ailsa craig (Wanless 1979).

| Year | Age Class | Feb | Mar | Apr | May | Jun | Jul | Aug | Sep | Oct | Totals |
| --- | --- | --- | --- | --- | --- | --- | --- | --- | --- | --- | --- |
| 1974 | Ad. | 0 | 14 | 9 | 7 | 16 | 26 | 23 | 13 | 0 | 108 |
|  | Imm. | 0 | 0 | 0 | 0 | 0 | 2 | 2 | 1 | 0 | 5 |
| 1975 | Ad. | 0 | 5 | 22 | 17 | 20 | 28 | 21 | 19 | 0 | 132 |
|  | Imm. | 0 | 0 | 1 | 0 | 7 | 4 | 1 | 2 | 0 | 15 |
| 1976 | Ad. | 9 | 10 | 17 | 13 | 14 | 18 | 25 | 20 | 1 | 127 |
|  | Imm. | 0 | 0 | 0 | 0 | 2 | 1 | 1 | 0 | 0 | 4 |
| Totals | Ad. | 9 | 29 | 48 | 37 | 50 | 72 | 69 | 52 | 1 | 367 |
|  | Imm. | 0 | 0 | 1 | 0 | 9 | 7 | 4 | 3 | 0 | 24 |

### Statistical modelling of crash probability

To evaluate if environmental conditions impacted gannet crashes at Ailsa Craig (objective 1), we obtained wind estimates from ERA5, a weather reanalysis tool that allows both forecasting and hind-casting of wind conditions. Specifically, hourly wind speed and direction were derived from the zonal (u) and meridional (v) wind components for a 0.25° × 0.25° grid cell centred on the breeding colony (55.254° N, 5.121° W) (Hersbach et al. 2023). As carcass counts were conducted monthly, we aggregated wind speed and direction data to monthly proportions.

Horizontal visibility data (fog) were obtained from two nearby coastal airports, Prestwick and Cambeltown (previously known as the Machrihanish RAF base). Data were obtained from the Met Office MIDAS database archived at CEDA (Centre for Environmental Data Analysis) (Met Office 2012). Visibility data were available at hourly or 3 hourly intervals during the study period. These data were summarised to give the number of days with fog per study month, with a day being defined as foggy if it included at least one measurement with visibility < 1000 m.

We conducted preliminary analyses to examine whether wind speed influenced crash mortality. To address the mismatch between monthly carcass counts and hourly wind estimates we grouped hourly wind speeds into five categories (0–2, 2–4, 4–6, 6–8, and >8 m s^-1^), providing the number of hours each month that fell within each wind class. These categories were included as predictors in models of monthly crash probability, but wind speed was not statistically significant. Based on this result, we focussed only on the two extremes of wind conditions in subsequent analysis, calm (<2 m s^-1^) and high (>8 m s^-1^). This tested for potential effects of unusually low or high wind speeds on crash probability.

There were two modal wind directions during the study period: north westerlies and south easterlies. We therefore categorised wind direction into two broad directional bins, taking northwest (NW) and southeast (SE) as the midpoint of each bin, each of which encompassed 180°. We ruled out collinearity with wind direction, fog, bird count and wind speed, by testing them against each other using the ggpairs in the GGally package (Schloerke et al. 2025).

Monthly surveys of colony size throughout the study period (Wanless 1979) revealed notable fluctuations through the season. We therefore modelled the monthly crash probability as the number of crashes divided by the number of birds at the colony per month. Wind speed, direction and the number of foggy days were included as fixed effects. We also included month, to account for the possibility that flight behaviour changes through the season, and colony size, to test whether crash probability was influenced by crowding at the colony. Binomial GLMs were initially applied, as the response variable was binary, with 0 indicating a bird that did not crash and 1 indicating a bird that did. However, post-hoc diagnostic tests revealed evidence of overdispersion in the data. Therefore, final models used a quasibinomial error distribution. Statistical models were constructed and visualised using R v4.4.1 (R Core Team 2024). We used the packages ggplot2 and visreg (Breheny and Burchett 2017) for visualising results.

We excluded October from the statistical analysis as while surveys were conducted from February to October, October bird counts were consistently low, with a mean of 5,200 across all years (figure S1) (Wanless 1979). Furthermore, gannets may be completely gone from colonies by early to mid October depending on the year (Nelson 2010). Whilst the use of crashes per bird at the colony adjusts for low bird counts, including months when birds leave the colony early would introduce error in the model as the proportion of wind speeds and direction is estimated per month.

### Modelling airflows over Ailsa Craig

Airflow characteristics over Ailsa Craig were modelled using the computational fluid dynamics (CFD) package OpenFOAM (openfoam.com version v2412). This has been validated over steep islands (Bechmann et al. 2011) and is widely used for modelling flows in the atmospheric boundary layer. Our approach and boundary conditions followed Lempidakis et al (2022). In summary, the initial coarse model domain was 7000 x 6000 x 1200 m, with a horizontal and vertical resolution of 40 m. The bottom boundary represented the surface of the island which was taken from a DEM of Ailsa Craig with 5 m resolution (‘Digimap’, n.d.). After establishing the initial mesh, the tool snappyHexMesh in OpenFOAM was used to incorporate the DEM, refining initial mesh cells close to the surface up to 3 times, adding 3 finer resolution layers near the surface and performing standard mesh quality tests and corrections (https://cfd.direct/openfoam/user-guide/v6-snappyhexmesh/). This resulted in a finer mesh resolution close to the surface of 5 m in the horizontal and 1 m in the vertical. Simulations were complete once convergence had been achieved using a steady-state incompressible solver with a k-ε turbulence closure scheme (using standard settings) (Lempidakis et al. 2022).

Simulations of the airflows over Ailsa Craig were run for the two modal wind directions; southerly and north-westerly. In both scenarios the at-sea wind speed was 10 m s^-1^ at 20 m height. The horizontal and vertical wind components (mean U and W respectively), and turbulent kinetic energy (TKE, a measure of the absolute wind unsteadiness) were extracted 2 m normal to the ground surface in order to visualise the conditions that gannets experience when flying close to ground.

### Matrix population model construction

Northern gannets are long-lived seabirds that typically recruit in their fifth summer. We therefore divided their life cycle into five stages: fledgling (1^st^ year), juvenile 2nd year, juvenile 3rd year, juvenile 4th year, and adult. We incorporated this age-structured framework into Matric Population Models, MPMs (figure 1). We constructed and visualised all MPMs in R v4.4.1 (R Core Team 2024), using the tidyr (Wickham, Vaughan, et al. 2025) and dplyr (Wickham, François, et al. 2025) packages for matrix construction, and popbio (Stubben and Milligan 2007) and ggplot2 (Wickham 2016) for time series visualisation.

**Figure 1.**
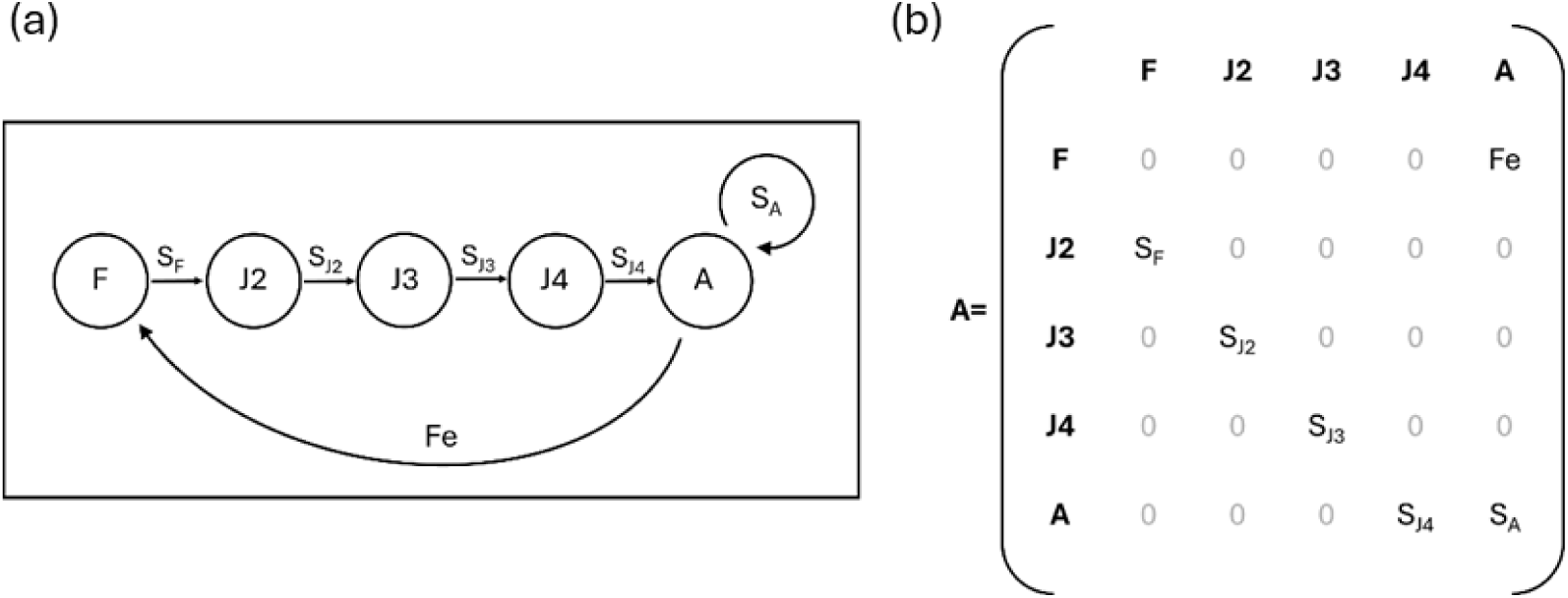
(a) A simplified diagram of the six stage Leslie style matrix population model (MPM) used where, C = chick, F = fledgling, J2-J4 = juvenile in their 2^nd^ to 4^th^ year respectively and A = adult. Arrows pointing right indicate a life stage transition. S_C_ = chick survival probability, S_F_ = fledgling survival probability, S_J2_ – S_J4_ = 2^nd^ to 4^th^ year juvenile survival probability and S_A_ = adult survival probability. Fe = adult fecundity. (b) Generic MPM matrix (X) with transition probabilities (S) of chicks (C), fledglings (F), Juveniles 2-4 (J2-4) and adults (A). Fecundity is represented as Fe in the top right of the matrix.

We obtained life stage transition rates for gannets from Wanless et al. (2006), who estimated them using ring recovery data, analysed with multinomial stochastic models (White and Burnham 1999) (table S1). The dataset included recoveries from 44,582 chicks ringed between 1959 and 2002 across 10 of the 19 UK gannet colonies and 1,445 adult gannets ringed at three major colonies. Bass Rock, the UK’s largest gannet colony, showed significantly different transition rates compared to all other colonies and was therefore treated separately. Since the present analysis focuses on Ailsa Craig, we took transition rates from the ‘other colonies’ category, of which Ailsa Craig made up the greatest proportion.

Fecundity refers to the reproductive output of individuals or populations, often measured as the number of offspring produced per individual (Bradshaw and McMahon 2008). In this analysis, we defined fecundity as the transition from adult to fledgling, representing the successful production of a fledgling by a breeding female. Gannet hatching success was estimated at 0.817 based on Wanless et al. (2006). We followed the MPM convention of halving this value to reflect only the female contribution, as gannets lay a single egg per breeding season and show no sex bias (Birdlife International 2018). We used the resulting fecundity value of 0.4085 as the adult-to-chick transition term in the MPMs. We later multiplied with the chick to fledgling transition probability. Although some individuals may breed before their fifth year (Nelson 1966), no quantitative estimates of early fecundity were available. Furthermore, sensitivity analyses indicated that including low-probability early breeding transitions had negligible effects on overall population growth. These transitions were therefore excluded from adjustment in the MPMs.

We took the probability of a chick successfully fledging as 0.737, based on estimates from Wanless and Harris (2003) and consistent with Wanless et al. (2006). We then multiplied this value with fecundity to give an accurate representation of the probability of a female gannet producing a successful fledgling. Although this transition occurs over several months within the breeding season, we treated it as a discrete annual stage to maintain consistency with the age-structured MPM framework. This simplification captures key biological timing, while allowing for integration of crash mortality in the MPM framework, and ultimately enabling me to adjust the fecundity in response to environmental conditions.

### Incorporating crash-induced mortality into matrix population models

We used MPMs to test the hypotheses that crashes influence gannet population dynamics. MPM transitions can be adjusted to reflect environmental pressures when carcasses or affected individuals are counted and accurately aged, allowing the proportion of mortality attributable to a given pressure to be quantified and incorporated into the model (Romanov and Masterov 2020). In this study, we used carcass count data from Wanless (1979) to estimate crash-related mortality, with adjustments applied to adult and chick-to-fledgling survival transitions. We created a hypothetical ‘no-crash’ scenario by calculating the proportion of annual mortality represented by crashed carcasses and removing this proportion from the relevant survival transitions. We integrated the significant outputs of the quasibinomial models predicting crash mortality into MPMs by adjusting transition rates according to the fitted relationships. This approach allowed us to evaluate both the direct demographic impact of crashes and the potential influence of shifting environmental drivers on crash-related mortality.

Wanless (1979), reported that most carcasses found at the base of the colony were adults. Across the three-year study period, only 24 out of 367 carcasses were classified as immature, averaging just eight individuals per year. Moreover, immature birds could not be reliably assigned to specific life stages (i.e. fledgling through fourth-year juvenile) due to uncertainty in gannet plumage and ageing. Given the low frequency of immature carcasses and the disproportionate influence of adult survival on population growth in long-lived species (Maestri et al. 2017; Romanov and Masterov 2020; Spencer and Janzen 2010), a pattern supported by sensitivity analyses, we did not adjust the immature survival transitions.

We adjusted the chick-to-fledgling transition under the assumption that if one parent died, the chick would fail. We then used this to adjust the adult to chick transition (0.4085) by multiplying them together. Northern gannets exhibit obligate biparental care, with both adults required for successful incubation, provisioning, and chick protection throughout the breeding season (Wojczulanis-Jakubas et al. 2018). The loss of a single parent typically results in nest failure due to starvation or exposure (Botha and Pistorius 2018). Because the timing of crashes relative to the breeding period could not be determined, and parent specific crash rates were unavailable, we assumed the full 5.4% annual crash mortality affected chick rearing.

### Incorporating stochastic variability into matrix population model projections

To improve the robustness of MPM projections, we incorporated stochastic variance into both the survey population and ‘no crash’ scenarios. We used Beta distributions to simulate variability as survival probabilities are bounded between 0 and 1. For the survey population projection, we used confidence intervals for gannet survival rates reported in Wanless et al. (Wanless et al. 2006) to derive shape parameters for a beta distribution using beta distribution equation (Johnson et al. 1995). Then, we used these parameters to simulate 1000 iterations of survival transitions in R studio, generating confidence ribbons around the survey population trajectory.

For the population projection under the ‘no crash’ scenario, we calculated confidence intervals from mean carcass counts reported over three years (Wanless 1979). We converted these to shape parameters for a beta distribution using the equations for beta distribution. We used simulated values to increase chick and adult survival 1000 times, with each combination run through the MPMs.

## Results

### Environmental drivers of crash mortality at Ailsa Craig

ERA5 reanalysis data revealed two modal wind directions at Ailsa Craig, with frequent winds from both NW and SE (figure 2). Our model revealed that gannet crash rates increased with the proportion of NW winds (Log odds = 1.62, SE = 0.57, z = 2.87, p = 0.009, binomial GLM), suggesting that wind direction plays a key role in mortality at the colony. McFadden’s pseudo-R² indicated that approximately 17.1% of the variation in carcass rates was explained by the model. Month was also a significant predictor of crash probability. However, the effect size was extremely low (table 2).

**Figure 2.**
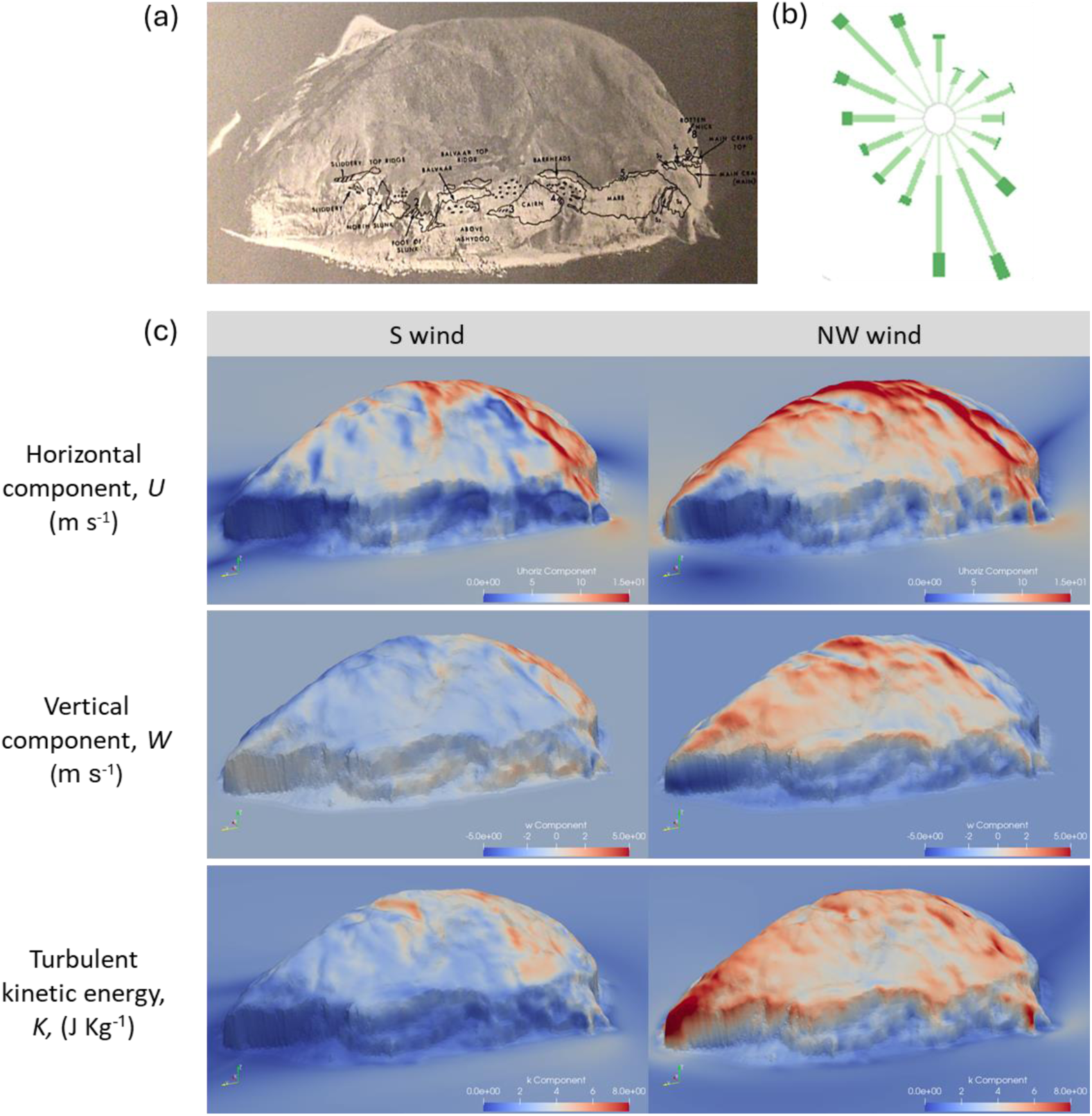
(a) Aerial photograph of Ailsa Craig taken from Wanless (1979) with gannet colonies outlined in black. (b) ERA5 wind directions for Ailsa Craig for the study period, showing 2 prevailing wind directions. (c) The airflows over Ailsa Craig as modelled in OpenFoam for southerly and north-westerly winds, where red colours show an increase in intensity of the horizontal component (top two panels), the vertical component (middle panels and the turbulent kinetic energy (bottom two panels).

**Table 2.**
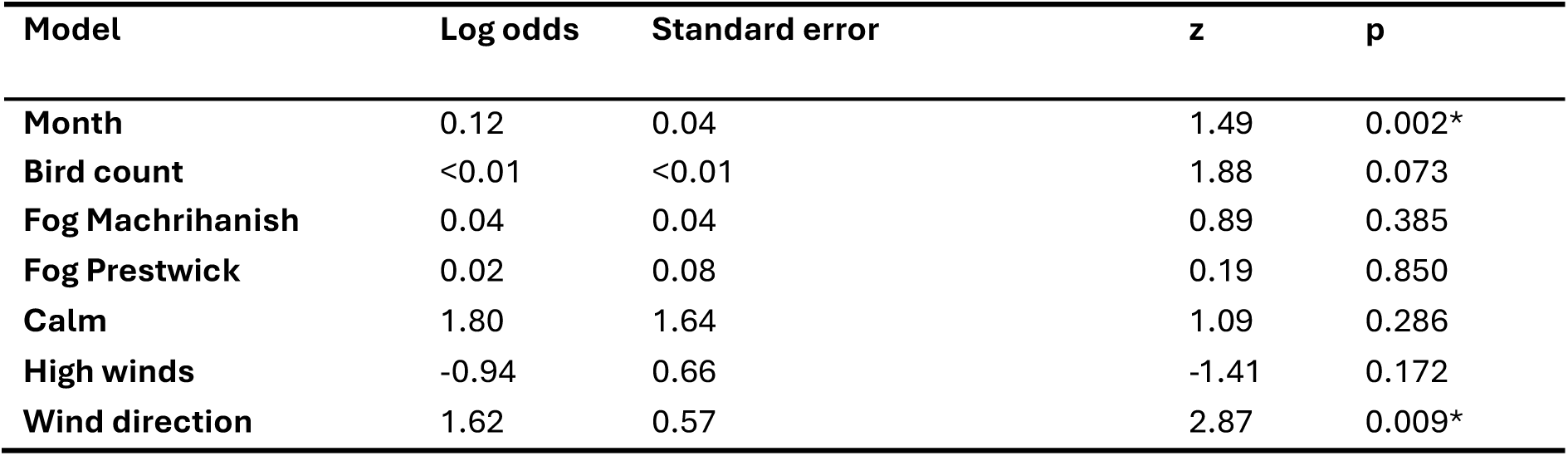
Results from independent generalized linear models (GLMs) with quasibinomial error families where crash probability was tested against environmental factors. Only month and wind direction emerged as significant results. Despite being significant, month had a very low effect size.

Pairwise correlation plots (figure S2) revealed a moderate positive correlation between bird count and month (r = 0.53). Bird count was also positively associated with calm wind conditions and negatively with high winds. Foggy days were uncorrelated with other variables, except for a weak association with calm conditions. NW wind proportion showed weak positive correlations with both month and bird count, indicating some seasonal trend but not enough to suggest problematic collinearity. Overall, there was little evidence to suggest that any other factors should be included when testing NW wind proportion against crash probability.

### Wind direction changes airflow characteristics along gannet breeding cliffs

OpenFoam CFD simulations show differences in all three airflow components around Ailsa Craig in the two wind directions (figure 2). Focusing on the characteristics along the breeding cliffs, the horizontal component showed the least change between the two wind directions, with only a slight increase in NW winds. The vertical flow component showed a more marked change with an increase in the strength of downdrafts in NW winds. The greatest change between wind directions was seen in TKE, which was substantially higher in NW winds all along the breeding cliffs.

### Effects of crash mortality on population growth rates

Our first MPM was run with the survey data to check that the MPM provided a realistic prediction of known colony growth rates. This model predicted that the gannet population on Ailsa Craig was experiencing steady growth, with a dominant eigenvalue (λ) estimated at 1.008 (table 3). This was reasonably well-aligned with the census data from Ailsa Craig, which was used to calculate an estimated ‘true’ population growth rate (λ) of 1.020 from a negative binomial GLM (1.015-1.025 95% CIs) (figure S4).

**Table 3.**
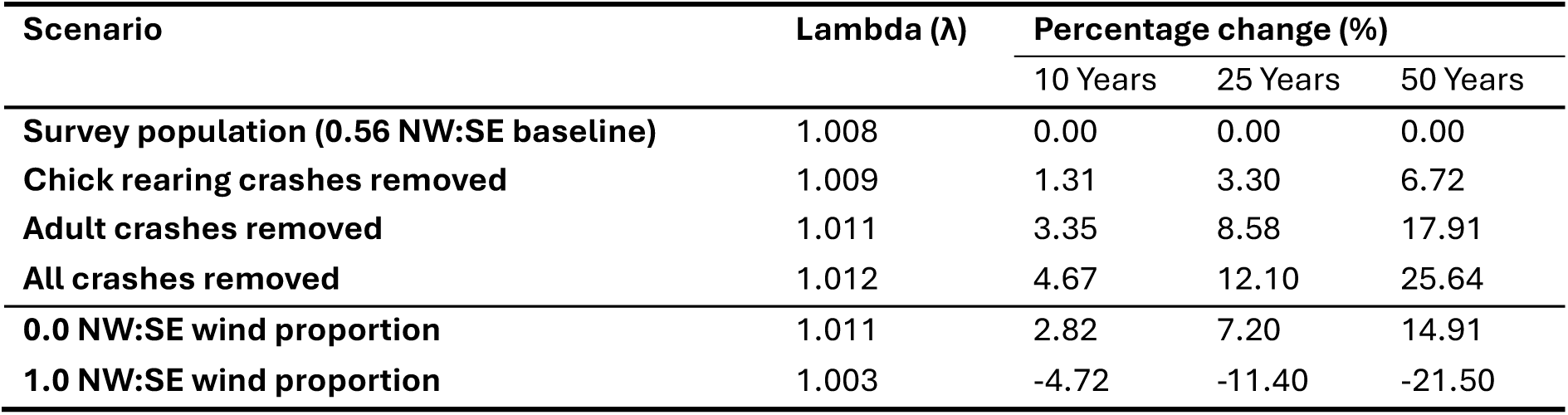
Predicted population growth rates for scenarios where adult crashes were included (survey population), only affected chicks (chick rearing crashes removed), only affected adults (adult crashes removed) and were excluded entirely (All crashes removed). Table also includes demographic predictions for 2 different wind scenarios where the proportion of north-westerly winds are fixed at 0.0 and 1.0. The dominant eigen values are given for each scenario, as well as the percentage changes from the survey population scenario over 10, 25 and 50 years.

We modelled a ‘no crash’ scenario where adult and chick-to-fledgling survival transitions were modified to remove this source of mortality. Adult crashes at the base of the colony accounted for an estimated 5.4% of annual adult mortality. This was used to adjust the adult survival transition from 0.922 to 0.9264 and the chick-to-fledgling transition from 0.737 to 0.752. This carried through to change fecundity from 0.301 to 0.307 (table S2).

The modified ‘no crash’ MPMs indicated that, over a 50-year projection period, the elimination of crash mortality is associated with a 25.6% increase in adult population at Ailsa Craig, corresponding to an increase from 41,272 to 51,855 adult birds (table 3). The reduction in adult mortality contributed most strongly to the observed population growth, as also shown in the sensitivity analysis of the original model. Decreases in chick-to-fledgling mortality had a more limited demographic effect, resulting in a 6.7% increase or around 2,800 additional adult gannets (table 3; figure S5).

Adult survival emerged as the most influential demographic rate. It had the highest sensitivity value (0.71), indicating that small absolute changes in survival would strongly affect population growth and have the greatest impact on λ. In contrast, adult fecundity had a lower sensitivity (0.19), suggesting smaller absolute influence. All other transitions had sensitivity values lower than that of adult fecundity.

### Demographic consequences of shifting wind regimes

Integrating the output of the quasibinomial GLM with the MPMs demonstrated how increasing the proportion of NW winds would have a negative impact on gannet population size at Ailsa Craig. In a theoretical scenario where all winds are NW, and all other factors, including colony location, remain the same, the population is projected to show a 21.50% reduction compared to the current wind scenario (0.564 proportion NW winds). Conversely, if all winds shifted to SE, the population is expected to increase by 14.91% compared to survey population. Together, these scenarios illustrate the possible envelope of population outcomes under future wind regime shifts (figure 4; table 3).

**Figure 4.**
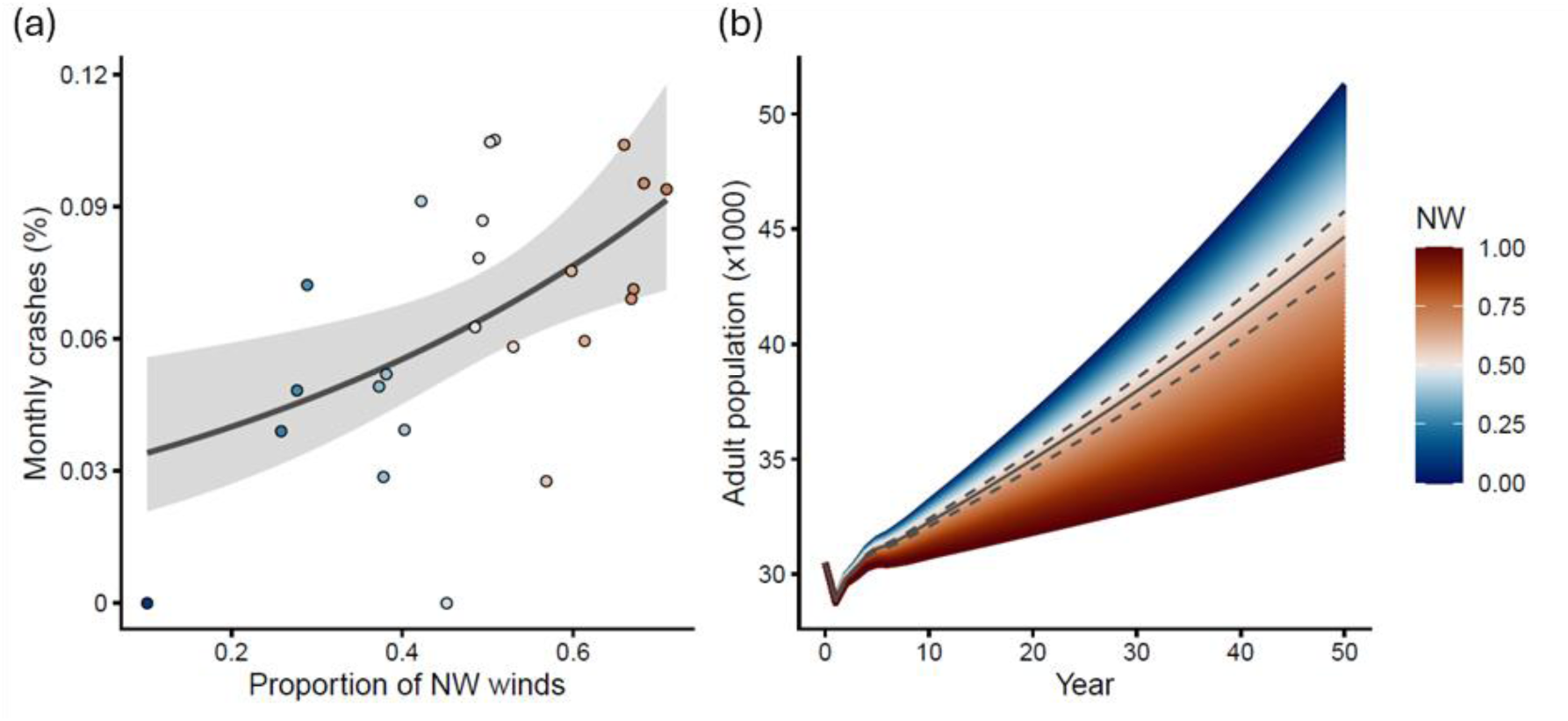
(a) The monthly crash probability, calculated from gannet carcass and bird count data (Wanless 1979), as a function of the proportion of north westerly (NW) winds. The fitted line represents a generalized linear model (GLM) with a quasibinomial error structure, including 95% confidence intervals. Points are coloured according to the monthly proportion of NW winds, using the same colour scale as in the right panel. (b) Matrix population model projections over 50 years under a range of hypothetical NW:SE proportions. Each coloured trajectory assumes a constant long-term proportion of NW winds between 0 (blue) and 1 (red), with colour indicating the NW proportion. The solid black line shows the projection for the observed mean NW wind proportion at Ailsa Craig (0.56) during the study period, and dashed lines show the 95% prediction interval of wind conditions over the past 50 years at Ailsa Craig. The model incorporates survival adjustments derived from the GLM in the left panel, simulating the demographic consequences of shifts in prevailing wind direction.

## Discussion

Adult gannets have high annual survival rates and are long-lived, making their populations particularly sensitive to perturbations in adult survival (Spencer and Janzen 2010; Wanless et al. 2006). In recent years, anthropogenic sources of mortality in adult gannets have become increasingly well documented, including fisheries bycatch (Araújo et al. 2022; Calado et al. 2021), collisions with wind turbines (Peschko et al. 2021; Pollock et al. 2021), and, most notably, outbreaks of highly pathogenic avian influenza (HPAI) (Giralt Paradell et al. 2023; Lane et al. 2024).

By contrast, non-anthropogenic pressures remain difficult to quantify, particularly given the species’ pelagic lifestyle. Here, we investigate crash-based mortality as a novel, non-anthropogenic pressure affecting the Ailsa Craig colony. We show that these crashes can substantially impact the local population, potentially reducing adult numbers by up to 21.5% over a 50-year period. Furthermore, we demonstrate that crash frequency is linked to wind direction, with higher proportions of north-westerly winds associated with increased mortality. To our knowledge, this is the first study to demonstrate that crash-based mortality can have a measurable demographic impact on population-level processes.

Crash-based mortality did not increase with the number of foggy days, indicating that collisions were unlikely to result from impaired perception of distance to the ground. Instead, crash rate was predicted by wind direction, with crashes increasing with the frequency of NW winds. Computational fluid dynamics (CFD) models predict that downdrafts are stronger along the breeding cliffs in NW winds, compared to southerlies, which are the other modal wind direction. Downdrafts are likely to be challenging for birds flying close to, and landing on, cliffs. Nonetheless, it is also conceivable that gannets could find solutions to this, given that NW winds are relatively common, and alter their approach strategy accordingly. We propose that the deciding factor that will push conditions beyond those that birds can respond to, is the turbulence associated with the downdrafts. In support of this, the most marked change brought by NW winds is an increase in the turbulent kinetic energy at the breeding cliffs in NW winds. Birds flying near the colony will therefore experience downdrafts with strong and unpredictable eddies that could cause temporary loss of flight control, in an area where they have very limited space to recover.

Against our expectations, wind speed did not predict gannet crashes. We predicted that calm winds could be more risky during landing as birds are typically use headwinds to reduce their groundspeed and landing impact. Wind can also generate updrafts that could improve control in the final phase of the landing. However, our analyses found no evidence that low wind speeds were more risky. Neither was there evidence that risks increased in strong winds. Auks abort their landing attempts in high winds and it was hypothesised this was due to the difficulty of responding to strong gusts (Shepard et al. 2019). While we found no evidence for wind speed affecting crash rate, it is difficult to rule wind speed out completely as crash rates were calculated from a single monthly count and predicted using aggregated monthly wind data. It may be that wind speed is unimportant relative to wind direction and turbulence, it could also be that high and low winds have their own attendant risks that will need further data to disentangle.

Removing crash-related mortality resulted in a predicted increase in population growth rate from λ = 1.008 to λ = 1.013. This change may appear small, but it translates to a 25.6% increase in adult numbers over a 50-year period (table 3), or 10,583 more breeding birds, compared to a population with current levels of crash-based mortality. The level of annual adult mortality (5.4%) is comparable to losses from fisheries bycatch (Le Bot et al. 2019) and greater than an extreme weather year that led to prey shortages and mass breeding failure at colonies in the western Atlantic (Montevecchi et al. 2021). Therefore, while levels of mortality do not approach those associated with recent HPAI outbreaks, chronic crash-related mortality may still impose significant long-term demographic pressure on gannet populations (Camphuysen and Gear 2022; Lane et al. 2024).

The actual impact of crash-based mortality on population growth will depend on the wind conditions, and whether they change through time. The yearly proportion of NW winds at Ailsa Craig has been between 48% and 65% for several decades with no apparent trend. Prevailing westerlies are a feature of the north eastern Atlantic (Trigo et al. 2008) and while surface wind patterns could change (Robson et al. 2018; 2016), a complete shift to NW or Southerly winds at Ailsa Craig seems unlikely. Nonetheless, scenarios where either wind direction dominates result in projections that differ by ∼20,000 individuals after 50 years (all other factors being equal). This illustrates the full envelope of demographic outcomes that crashes could have at our study site, depending on the wind regime, and more generally, highlights the importance of wind direction on risk.

Crash-based mortality is likely to affect the demography of other populations, beyond gannets breeding on Ailsa Craig. This may be particularly significant for the grey-headed albatross population studied by Schoombie et al. (2023), as crashes account for an estimated 11% of annual adult mortality – over twice that recorded for gannets on Ailsa. Furthermore, natural adult survival in grey-headed albatrosses is higher than in northern gannets (0.951 compared to 0.922, Converse et al. 2009, Wanless et al. 2006). The sensitivity of demographic rates to changes in adult mortality therefore suggest that the consequences of crashes in these albatrosses could be substantially greater than predicted for gannets breeding on Aisla Graig.

The two colonies where regular crashes have been documented (Ailsa Craig and grey-headed ridge on Marion Island (Schoombie et al. 2023)) are unusual, but not unique in their topography as the breeding cliffs do not drop directly into the sea. Fatalities may be therefore be more likely if birds that lose flight control crash into the ground below. Accidents could still occur at other colonies, as a loss of flight control could lead to uncontrolled collision with the vertical cliff face or landings on the sea. Overall, it is difficult to assess how topography affects the probability or severity of accidents, but, importantly, only fatal crashes onto land are likely to be detected, as fatal crashes at sea or non-fatal crashes almost certainly go undocumented.

The risk of flight accidents should vary according to the combined effects of wind conditions, topography, and flight morphology (Shepard et al. 2019). Large mass should predispose species to a greater risk of accident when operating close to land as landing forces will increase with mass and the deceleration on impact. Larger flying animals also have lower available power (Pennycuick 2008), which could affect their ability to recover flight speed or height after a loss of control. Finally, high aspect-ratio wings should be more prone to fracture, as they experience greater torque than short wings if they make contact with a solid substrate. The probability and severity of accidents may therefore be particularly high in seabirds, which are typified by large mass and high-aspect ratio wings. Understanding how risk varies with morphology is a key priority given the growing footprint of anthropogenic infrastructure both on land and at sea, including the expansion of offshore wind developments, which combine physical obstacles with highly turbulent flow (Shepard 2025).

Overall, we identify wind-induced crashes as a substantial and previously overlooked source of adult mortality in northern gannets, a long-lived species whose population dynamics are highly sensitive to changes in adult survival. The MPMs suggest that this subtle, chronic, environmental pressure may play a key role in the population trajectory of the species at Ailsa Craig and potentially other colonies. Understanding how wind impacts seabirds is increasingly important as climate change continues to alter wind regimes globally (Pryor et al. 2012). The role of wind speed is generally well-understood through its impact on flight speed, flight costs and prey availability (Lewis et al. 2015; Pennycuick et al. 2013; Shepard et al. 2019), but much less is known about the importance of wind direction. Our results demonstrate that wind direction can influence the risk of mortality at the colony, suggesting a novel mechanism by which birds may be affected by changes in surface wind directions. While this study used data collected at Aisla Craig, risks relating to flight control should act as a selective pressure affecting seabirds more generally (KleinHeerenbrink et al. 2022; Lempidakis et al. 2022; Shepard et al. 2019). The risks of crashing may therefore already be shaping the spatial structure, survival, and evolutionary trajectories of cliff-nesting species in ways that remain largely undocumented.

## Acknowledgements

We are grateful to Emmanouil Lempidakis for his help with some exploratory modelling. We also thank Miguel Lurgi for stimulating discussions, and Prof. Owen Jones and Dr Kayleigh Rose for their helpful comments and feedback on Stefan Barnett’s thesis.

