## Supplementary material for "Flight crashes have demographic consequences in a long-lived seabird"

**Supplementary methods**

**Integrating wind-driven crash risk into matrix population models**

Wind was the main predictor of crash rate (see Results). Coefficients from the quasibinomial GLM were integrated with the MPMs to investigate how different wind conditions could influence population growth rates. The probability of crashing was calculated for a range of NW:SE wind proportions, using the equation for the GLM. Monthly crash probabilities were compounded across the 8 months when gannets were present at the colony to obtain yearly probabilities. This was calculated as one minus the probability of no crash occurring raised to the eighth power. To align these values with the mean NW:SE wind proportion between 1974 and 2024, the annual survival estimate (0.922) was at a NW:SE wind proportion of 0.56. A complete equation for this calculation is shown in Equation 1. This produced wind adjusted annual survival rates, centred around the observed survival and mean wind direction proportion, which were used to modify transition probabilities in the MPMs (table S1).

*Equation 1*

$$s_{\text{adj}}\left( NW \right)=s_{\text{base}}\cdot\frac{\left( 1 - \frac{e^{\beta_{0}+\beta_{1} NW}}{1+e^{\beta_{0}+\beta_{1}NW}} \right)^{8}}{\left( 1 - \frac{e^{\beta_{0}+\beta_{1} NW_{ref}}}{1+e^{\beta_{0}+\beta_{1}NW_{ref}}} \right)^{8}}$$

***Equation S1.*** *formula used to adjust chick fledging transition probability where:* $s_{\text{adj}}\left( NW \right)$*= wind-adjusted annual adult survival at wind proportion NW,* $s_{\text{base}}$ *= survey population annual adult survival (fixed at 0.922 when NW = 0.56),* $\beta_{0}$ *and* $\beta_{1}$*= intercept and slope from the quasibinomial generalised linear model,* $NW$ *= proportions of wind from the 180° NW direction,* $NW_{ref}$ *= the mean NW proportion since 1974 (0.56) and* $\left( 1-p_{\text{crash}} \right)^{8}$*= the annual survival across 8 breeding months, where* $p_{\text{crash}}$ *is crash probability.*

To adjust chick fledging probability, it was assumed that chicks only survive to fledge if both parents survive the breeding season (Nelson 1966). For each year, the probability that both parents survive was calculated as the square of adult annual survival ($s^{2}$). As before, this survival probability was then rescaled relative to the mean NW:SE wind proportion between 1974 and 2024 (0.56). The relative scaling factor was used to adjust the survey population fledging probability from the unmodified MPMs ($p_{\text{fledge, base}}=0.737$), yielding an adjusted fledging transition probability that incorporates wind-driven parental crashes. A complete equation for this calculation is shown in Equation 2 (table S2).

*Equation*

$$f_{adj}\left( NW \right)=p_{fledge,base}\left( \frac{s_{adult}\left( NW \right)}{s_{adult}\left( NW_{ref} \right)} \right)^{2}$$

***Equation S2.*** *formula used to adjust chick fledging transition probability. Where:* $f_{\text{adj}}\left( NW \right)$ *= adjusted chick fledging transition at wind condition NW,* $p_{\text{fledge, base}}$*= Survey population fledging probability from the unmodified MPMs (0.737),* $s_{\text{adult}}\left( NW \right)$ *= adult annual survival under wind condition NW (from quasibinomial generalised linear model) and* $NW_{\text{ref}}$ *= adult annual survival at the reference NW proportion (0.56). The exponent 2 reflects the assumption that both parents must survive.*

**Supplementary tables**

***Table S1.*** *Survival probabilities and 95 % confidence intervals calculated by Wanless et al. (2006) for northern gannets in the UK between 1959 and 2002.*

| **Age class** | **All colonies** | **Bass Rock** | **Other colonies** |
| --- | --- | --- | --- |
| **1st year** | 0.424 (0.410–0.439) | 0.542 (0.516–0.567) | 0.420 (0.394–0.445) |
| **2nd year** | 0.829 (0.821–0.836) | 0.779 (0.765–0.793) | 0.852 (0.842–0.861) |
| **3rd year** | 0.891 (0.886–0.896) | 0.859 (0.848–0.869) | 0.908 (0.901–0.915) |
| **4th year** | 0.895 (0.889–0.900) | 0.863 (0.852–0.874) | 0.910 (0.903–0.917) |
| **Adult** | 0.919 (0.915–0.922) | 0.916 (0.910–0.922) | 0.922 (0.916-0.927) |

***Table S2.*** *Summary of transition probabilities used in matrix population models (MPMs) for gannets at Ailsa Craig. Including adjusted transitions based on carcass counts found at Ailsa Craig between 1994 and 1996 (Wanless 1979).*

| **Transition** | **Transition Probabilities** | |
| --- | --- | --- |
|  | **Survey population** | **‘No Crash’ Scenario** |
| **Chick – Fledgling** | 0.737 | 0.752 |
| **Fledgling – Y2 Juvenile** | 0.420 | 0.420 |
| **Y2 Juvenile – Y3 Juvenile** | 0.852 | 0.852 |
| **Y3 Juvenile – Y4 Juvenile** | 0.908 | 0.908 |
| **Y4 Juvenile – Adult** | 0.910 | 0.910 |
| **Adult** | 0.922 | 0.926 |
| **Fecundity** | 0.4085 | 0.4085 |

**Supplementary figures**

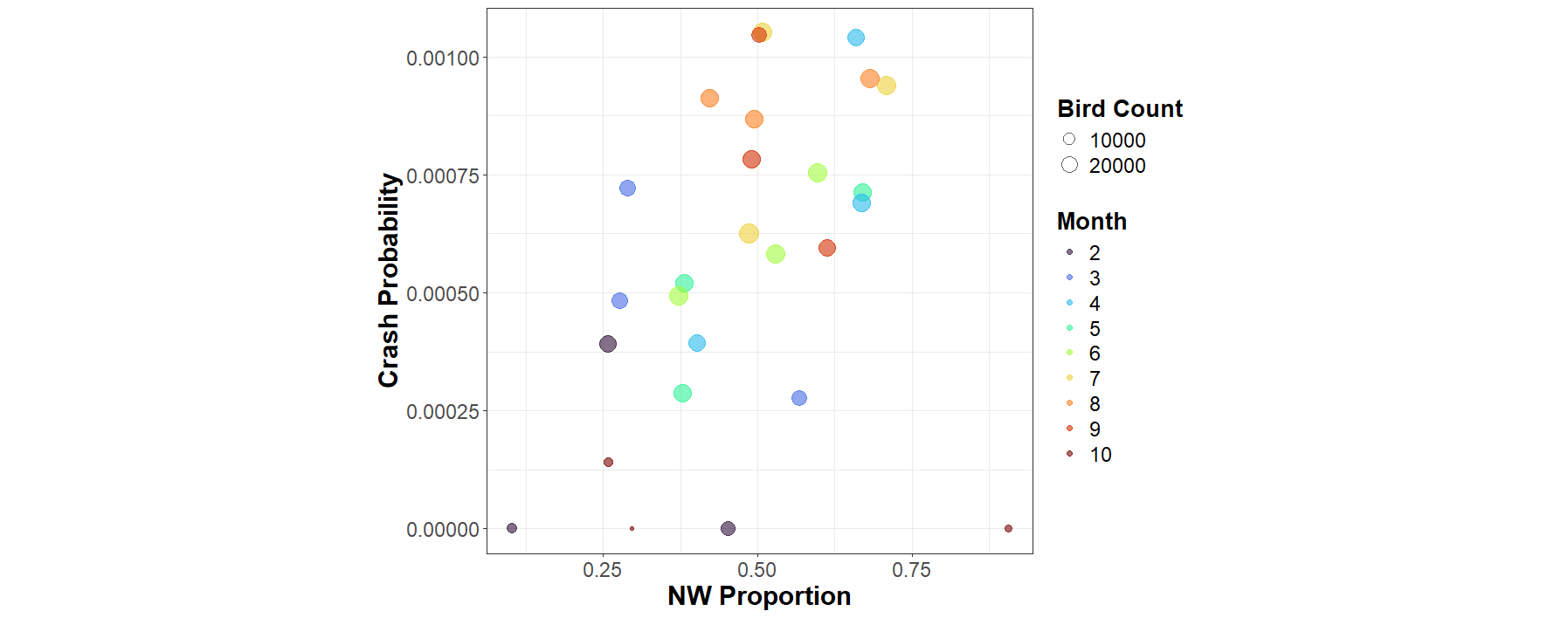

8

***Figure S1.*** *Monthly crash probability plotted against wind proportion (NW:SE) With points coloured by month with proportional sizes depending on the number of birds at the colony in that month and year. October points are circled*.

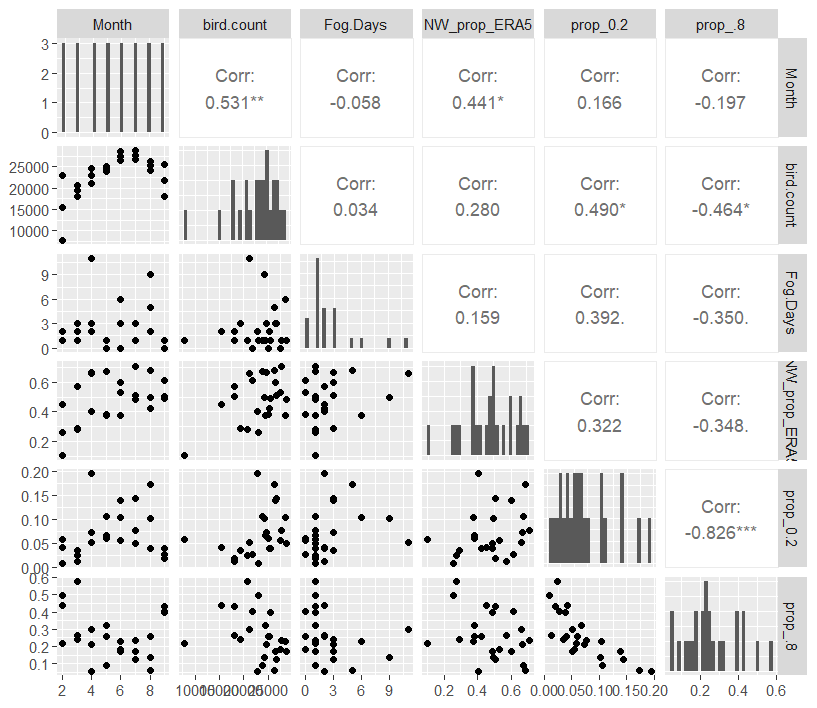

9

***Figure S2.*** *Testing for correlation between environmental factors at Ailsa Craig.*

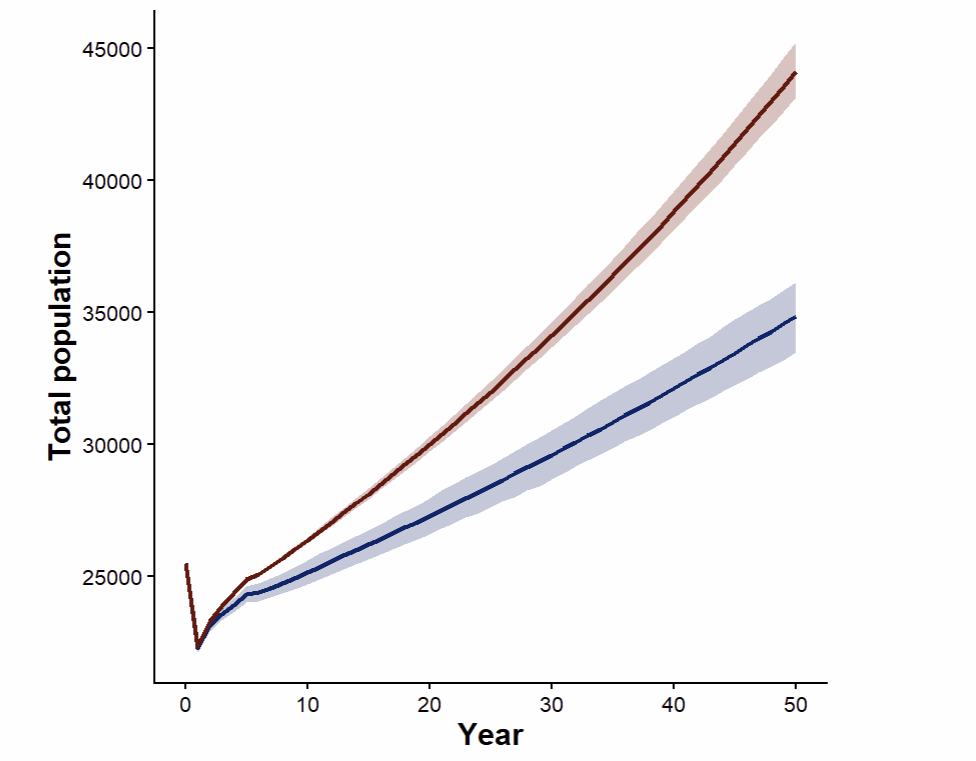

***Figure S3.*** *(a) Demographic predictions of the Ailsa Craig gannet population for scenarios with crash mortality (survey population), (blue) and without crash mortality (red). Lines include stochastic variation calculated from beta distribution of survival estimate confidence intervals (blue) and carcass count means (red).*

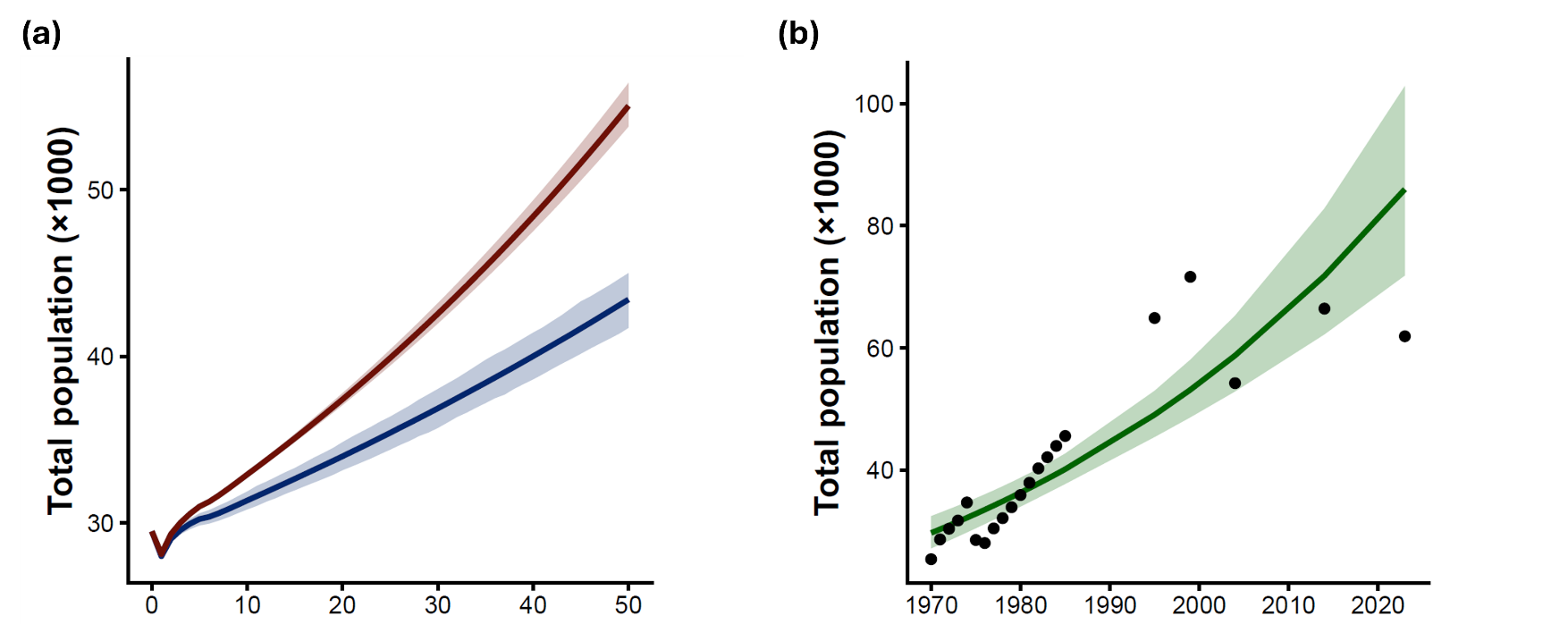
**Figure S4.** *The population trend at Ailsa Craig is indicated by black circles, which represent counts listed by the seabird monitoring program (SMP) and British trust for ornithology (BTO) for the period 1970 – 2023. Surveys were conducted yearly from 1970 – 1985 and then around once every 10 years from 1985 - present. The MPM projection for the Aisla population is indicated with a solid black line.*

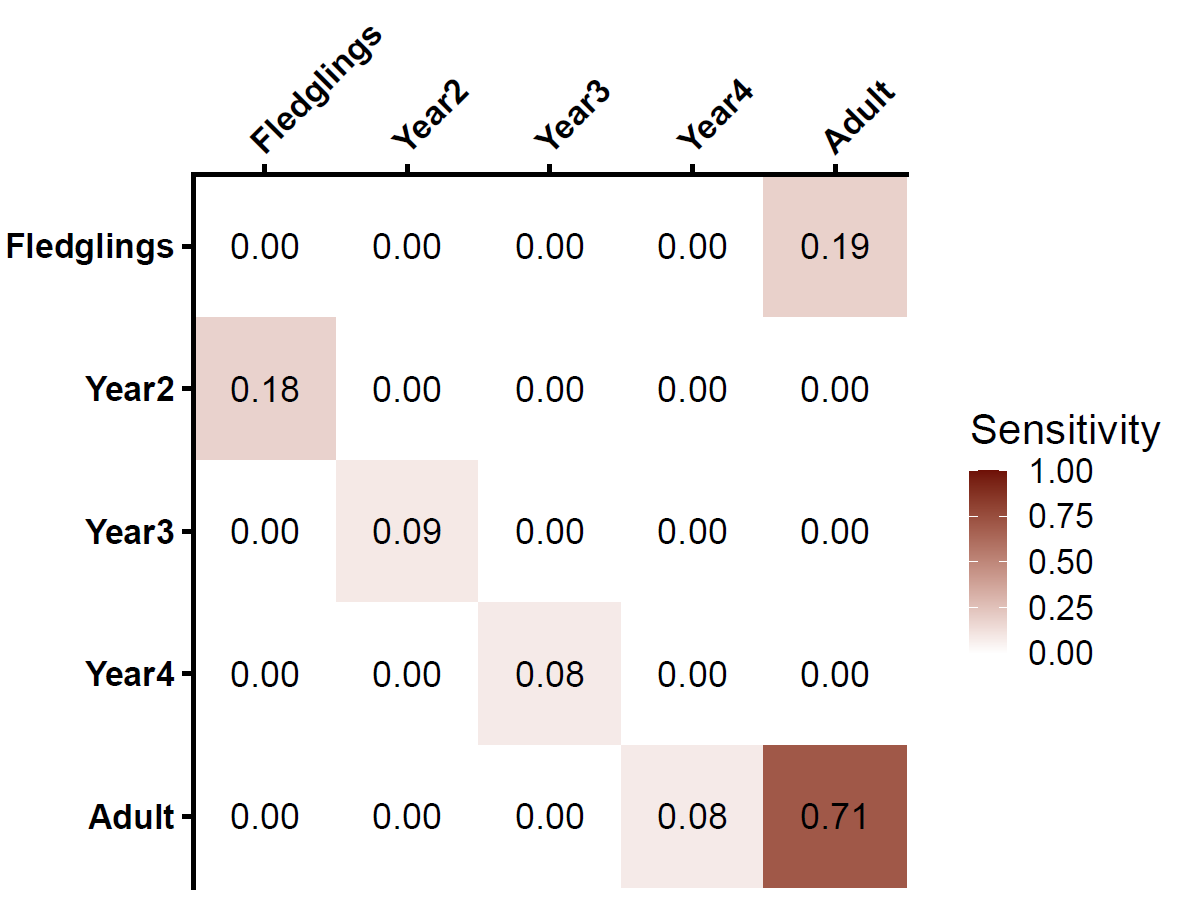

***Figure S5****. Sensitivity results from a matrix population model (MPM) growth rate to perturbations based on survival transitions in northern gannets at Ailsa Craig. Darker colours in both figures indicate greater values.*

5

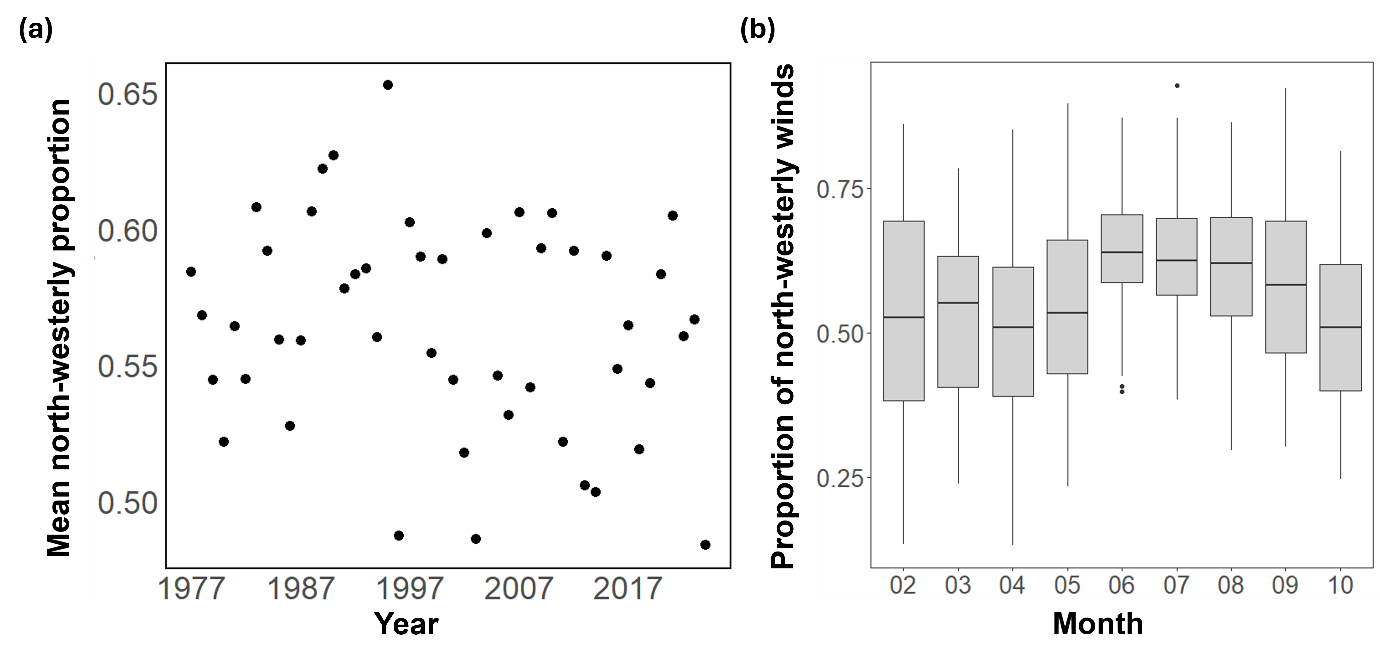

***Figure S6.*** *(a) Annual mean proportion of north-westerly (NW) winds at Ailsa Craig from 1977 to 2024, based on ERA5 wind reanalysis data. Each point represents the average NW wind proportion from February to October for a given year. Across the 48-year period, the proportion of NW winds per year ranged from 0.484 - 0.653 (mean of 0.564). This translates into predicted crash rates of 0.62% - 0.79% across a breeding season, using the coefficients from the quasibinomial GLM. No significant temporal trend was detected in the proportion of NW winds (linear model, SE = -0.001, p = 0.168), although the years used in our analyses (1974 to 1976) exhibited a slightly lower average NW wind proportion compared to the mean for the subsequent years. (b) Monthly variation in the proportion of NW winds across all years (February to October). An ANOVA revealed a significant effect of month on the proportion of NW winds throughout the year (F = 5.395, df = 8, 423, p < 0.001), indicating a seasonal pattern in wind directionality. NW winds were most frequent in June and July, while October had the lowest NW wind proportions. Boxes show interquartile ranges (IQR) with medians. Whiskers represent an additional 1.5× IQR. Further outliers are shown as individual points.*

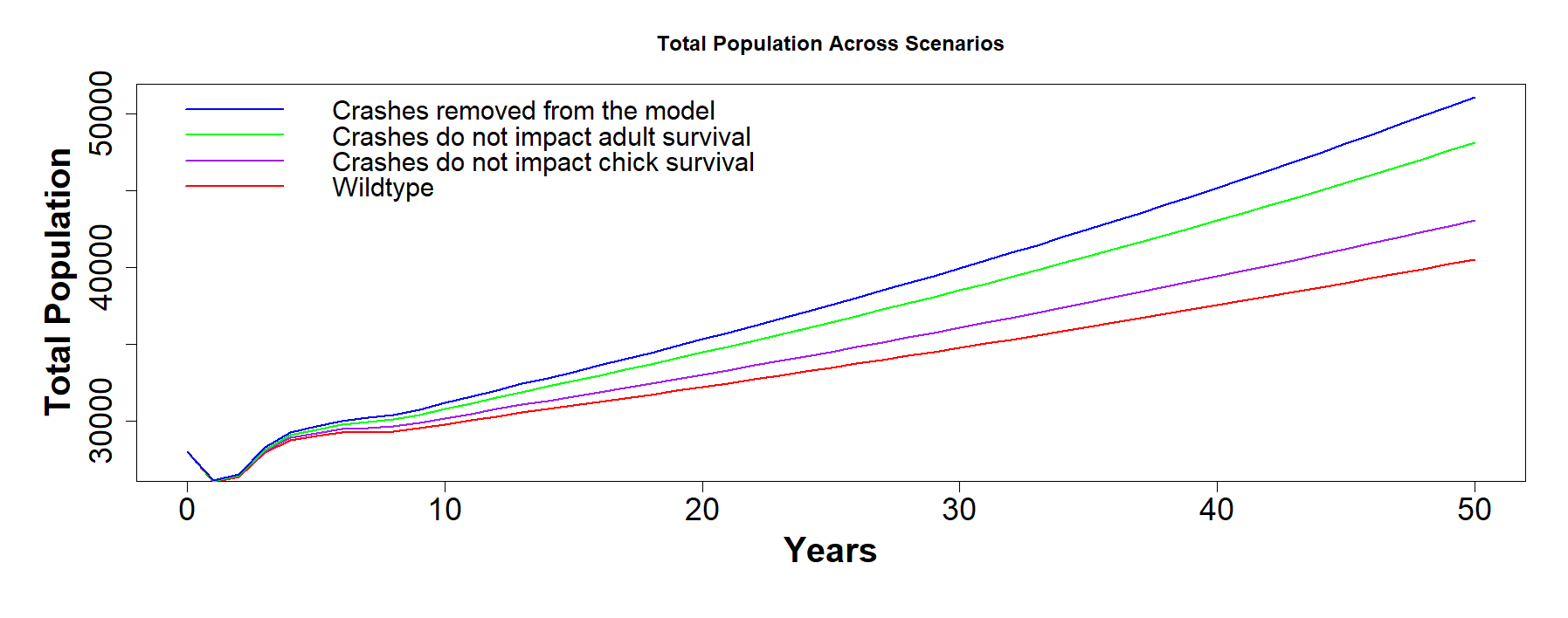

7

***Figure S7.*** *Projected population trajectories for the Ailsa Craig gannet colony under four demographic scenarios: the* *survey population* *model including crash-related mortality (red), crash mortality removed during chick-rearing only (purple), crash mortality removed from adult survival only (green), and all crash mortality removed (blue). These projections highlight the dominant influence of adult survival on long-term population dynamics, consistent with sensitivity analyses showing that changes to adult survival transitions have the greatest effect on population growth rate*

**References**

Wanless, Sarah. 1979. *Aspects of Population Dynamics and Breeding Ecology in the Gannet (Sula Bassana (l.)) of Ailsa Craig*. University of Aberdeen (United Kingdom).

Wanless, Sarah, Morten Frederiksen, Michael P. Harris, and Stephen N. Freeman. 2006. ‘Survival of Gannets Morus Bassanus in Britain and Ireland, 1959–2002’. *Bird Study* 53 (1): 79–85. https://doi.org/10.1080/00063650609461419.
